## Supplemental Information for "A comprehensive map of genetic interactions in childhood cancer reveals multiple underlying biological mechanisms"

### Supplementary information

*Table S1: Number of tumors and samples removed during filtering*

| DKFZ | Total | | WGS | | WES | |
|---|---|---|---|---|---|---|
| | Tumors | Samples | Tumors | Samples | Tumors | Samples |
| **In downloaded files** | **914** | **961** | **542** | **547** | **372** | **414** |
| Hypermutators | 7 | 7 | 5 | 5 | 2 | 2 |
| Single end sequencing samples | 10 | 10 | 0 | 0 | 10 | 10 |
| Relapses | 35 | 82 | 2 | 7 | 33 | 75 |
| **Remaining after sample filtering** | **862** | **862** | **535** | **535** | **327** | **327** |
| Samples without functional mutations | 33 | 33 | 12 | 12 | 21 | 21 |
| **Remaining after mutation filtering** | **829** | **829** | **523** | **523** | **306** | **306** |
| **TARGET** | **Total** | | **WGS** | | **WES** | |
| | Tumors | Samples | Tumors | Samples | Tumors | Samples |
| **In downloaded files** | **1,648** | **1,672** | **653** | **655** | **1,115** | **1,115** |
| Missing metadata | 3 | 3 | 1 | 1 | 2 | 2 |
| Hypermutators | 2 | 2 | 0 | 0 | 2 | 2 |
| **Remaining after sample filtering** | **1,643** | **1,667** | **652** | **654** | **1,111** | **1,111** |
| Samples without functional mutations | 11 | 11 | 5 | 5 | 6 | 6 |
| **Remaining after mutation filtering** | **1,632** | **1,656** | **647** | **649** | **1,105** | **1,105** |

*Note: after filtering one tumor per patient remains, so final tumor counts can be read as final patient counts.*

***Table S2: Candidate gene pairs in TARGET and DKFZ data set***

Table is separately available as excel file Table_S2.xlsx

### *Table S3: MLA scores of candidate gene pairs*

MLA scores for genes in candidate genetic interactions and absolute difference between their MLA scores (ΔMLA). 'Suspect' gene pairs are marked in grey and have at least one gene with MLA > 3 and have ΔMLA > 3 in case of mutual exclusivity or both MLA > 3 in co-occurring gene pairs. For the mutated genes involved in suspect gene pairs, the cancer sub type in which they are enriched are listed in parentheses. The table is sorted on ΔMLA for each data set.

| data set | cancer type | gene-1 (subtype) | gene-2 (subtype) | MLA-1 | MLA-2 | ΔMLA | suspect |
|---|---|---|---|---|---|---|---|
| **TARGET** | T-ALL | NOTCH1 | USP7 (TAL1) | 4.7 | -1.6 | 6.4 | x |
| | T-ALL | JAK3 (HOXA) | PTEN (TAL1) | 4.3 | -1.7 | 6.0 | x |
| | T-ALL | PHF6 (TLX3) | PTEN (TAL1) | 3.5 | -1.7 | 5.2 | x |
| | T-ALL | PHF6 (TLX3) | USP7 (TAL1) | 3.5 | -1.6 | 5.1 | x |
| | T-ALL | DNM2 | USP7 | 2.9 | -1.6 | 4.5 | - |
| | T-ALL | MAGI1 | NOTCH1 | 0.3 | 4.7 | 4.4 | x |
| | T-ALL | PTEN | WT1 | -1.7 | 2.5 | 4.3 | - |
| | T-ALL | USP7 | WT1 | -1.6 | 2.5 | 4.2 | - |
| | B-ALL | KRAS | TP53 | 3.0 | -0.8 | 3.8 | - |
| | T-ALL | FBXW7 | JAK3 (HOXA) | 1.1 | 4.3 | 3.2 | x |
| | T-ALL | FBXW7 | PTEN | 1.1 | -1.7 | 2.8 | - |
| | WT | DROSHA | TP53 | -1.1 | 1.7 | 2.8 | - |
| | T-ALL | JAK3 | STAT5B | 4.3 | 1.7 | 2.6 | - |
| | T-ALL | PHF6 | PIK3R1 | 3.5 | 1.0 | 2.5 | - |
| | AML | NRAS | WT1 | 2.7 | 0.5 | 2.1 | - |
| | AML | KIT | NRAS | 0.9 | 2.7 | 1.8 | - |
| | B-ALL | PXDN | ZNF582 | 4.1 | 2.8 | 1.3 | - |
| | B-ALL | CRLF2 | NRAS | 0.3 | 1.5 | 1.3 | - |
| | AML | CEBPA | CSF3R | 0.1 | 1.3 | 1.1 | - |
| | B-ALL | FLT3 | KRAS | 2.0 | 3.0 | 1.0 | - |
| | T-ALL | JAK1 (HOXA) | JAK3 (HOXA) | 3.5 | 4.3 | 0.8 | x |
| | T-ALL | NRAS | WT1 | 3.3 | 2.5 | 0.8 | - |
| | AML | FLT3 | IDH2 | 0.4 | 1.1 | 0.7 | - |
| | AML | FLT3 | KIT | 0.4 | 0.9 | 0.5 | - |
| | AML | CEBPA | WT1 | 0.1 | 0.5 | 0.4 | - |
| | AML | IDH2 | NPM1 | 1.1 | 0.7 | 0.4 | - |
| | AML | KIT | WT1 | 0.9 | 0.5 | 0.4 | - |
| | WT | CTNNB1 | EFCAB6 | -0.6 | -0.8 | 0.2 | - |
| **DKFZ** | HGG-K27M | ACVR1 | TP53 | -1.2 | 2.1 | 3.3 | - |
| | HGG-K27M | ACVR1 | H3F3A | -1.2 | 1.4 | 2.6 | - |
| | HGG-K27M | H3F3A | HIST1H3B | 1.4 | -1.1 | 2.5 | - |
| | HGG-other | ATRX | H3F3A | 0.7 | 1.9 | 1.2 | - |
| | MB-SHH | PTCH1 | SMO | -0.8 | 0.2 | 0.9 | - |
| | MB-WNT | DDX3X | KMT2D | 0.7 | 1.2 | 0.5 | - |
| | MB-SHH | KMT2D | PTCH1 | -0.9 | -0.8 | 0.1 | - |
| | HGG-K27M | ACVR1 | HIST1H3B | -1.2 | -1.1 | 0.1 | - |

### *Supplemental Figure S1: MLA distribution per TARGET cancer type*

Distribution of MLA scores for each TARGET cancer type in which candidate genetic interactions were found. Genes that are part of at least one candidate gene pair are shown with their MLA score and mutation frequency (secondary vertical axis).

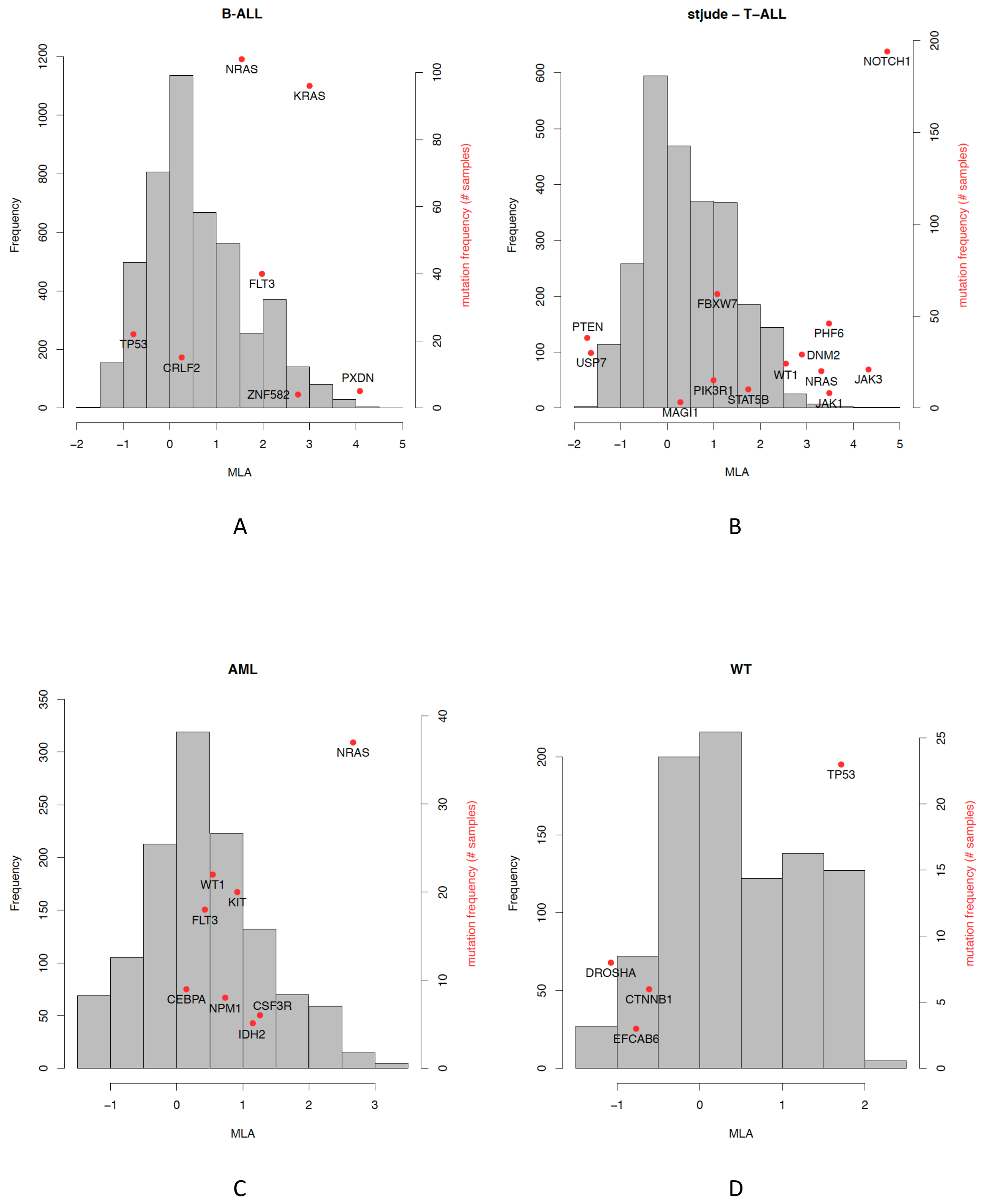

*Supplemental Figure S2: MLA distribution per DKFZ cancer type*

Distribution of MLA scores for each DKFZ cancer type in which candidate genetic interactions were found. Genes that are part of at least one candidate gene pair are shown with their MLA score and mutation frequency (secondary vertical axis).

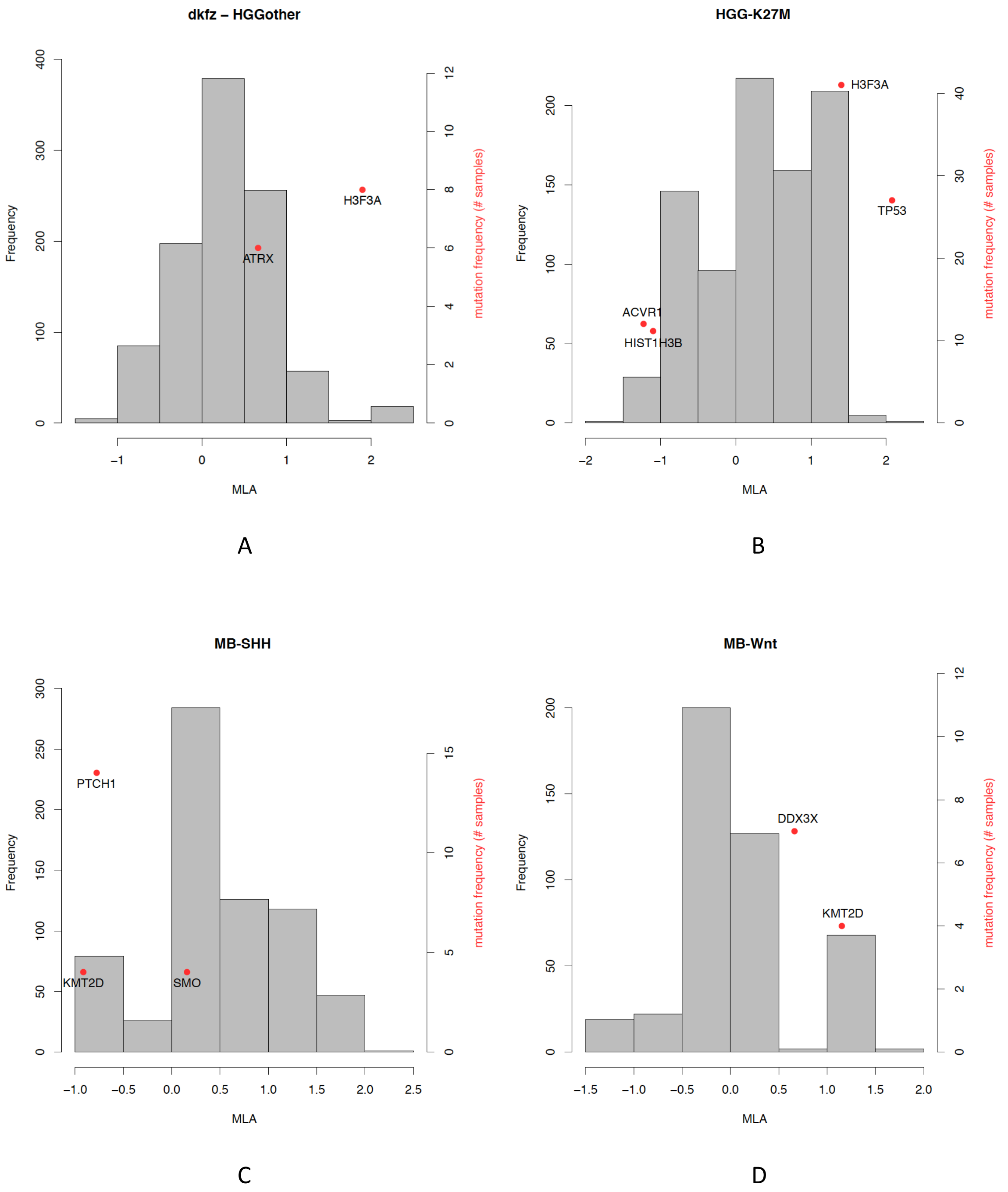

### *Supplemental Figure S3: Tumor load and mutation frequency of suspect gene pairs*

Mutation frequency heatmaps of both genes in the seven ‘suspect’ T-ALL candidate gene pairs (having at least one gene with MLA > 3 and having ΔMLA > 3 in case of mutual exclusivity or both genes having an MLA > 3 in co-occurrent pairs). Samples are ordered on tumor mutation load (TML).

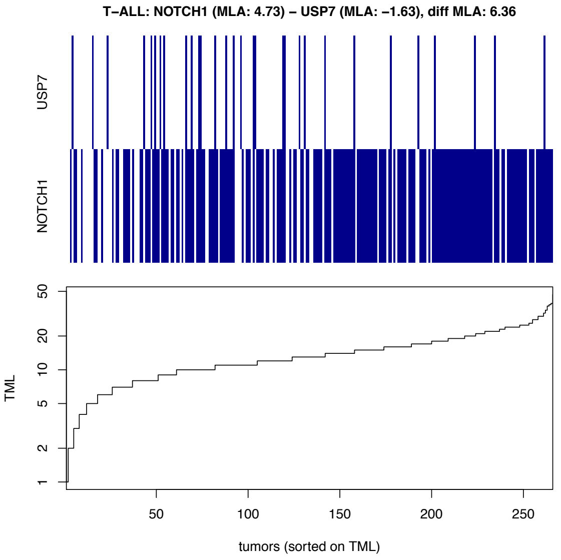

A

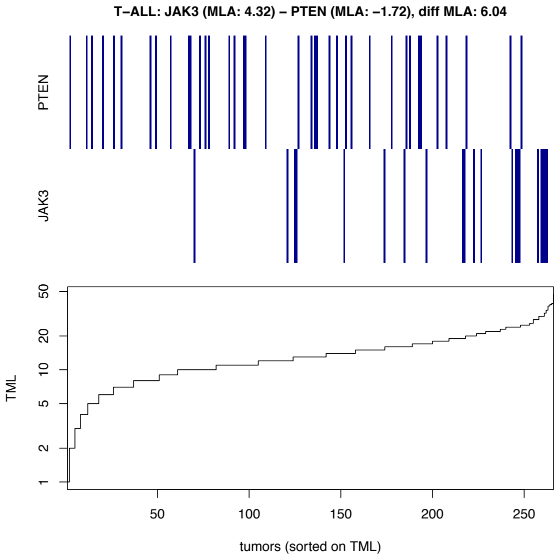

B

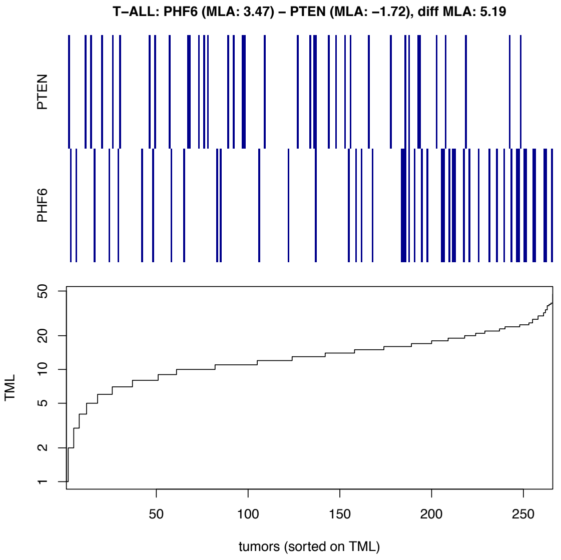

C

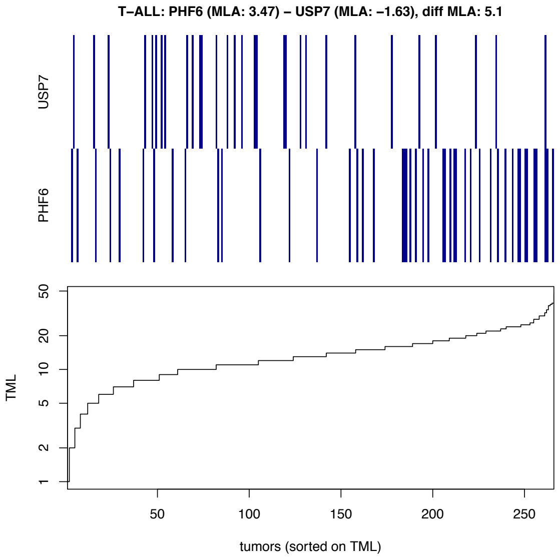

D

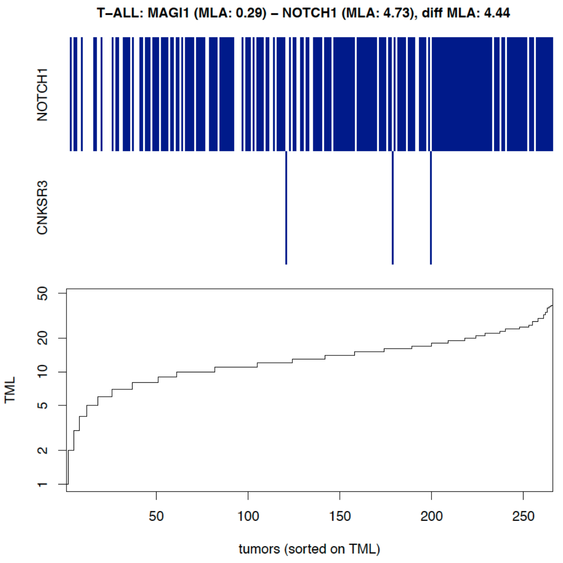
T–ALL: MAGI1 (MLA: 0.29) – NOTCH1 (MLA: 4.73), diff MLA: 4.44
NOTCH1
CNKSR3
TML
1
2
5
10
20
50
50
100
150
200
250
tumors (sorted on TML)

E

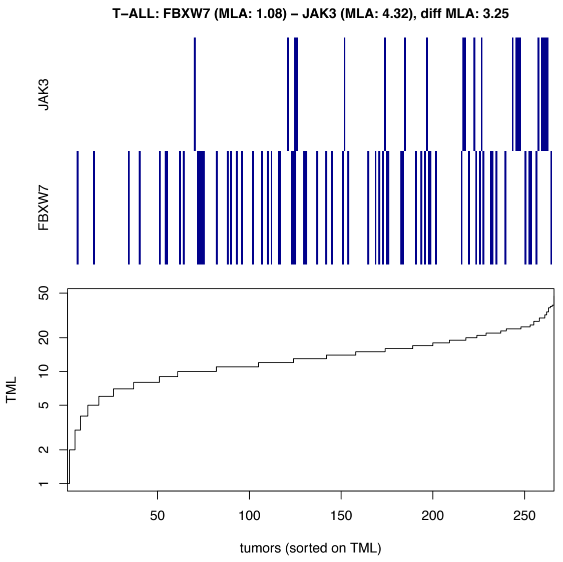
T–ALL: FBXW7 (MLA: 1.08) – JAK3 (MLA: 4.32), diff MLA: 3.25
JAK3
FBXW7
TML
1
2
5
10
20
50
50
100
150
200
250
tumors (sorted on TML)

F

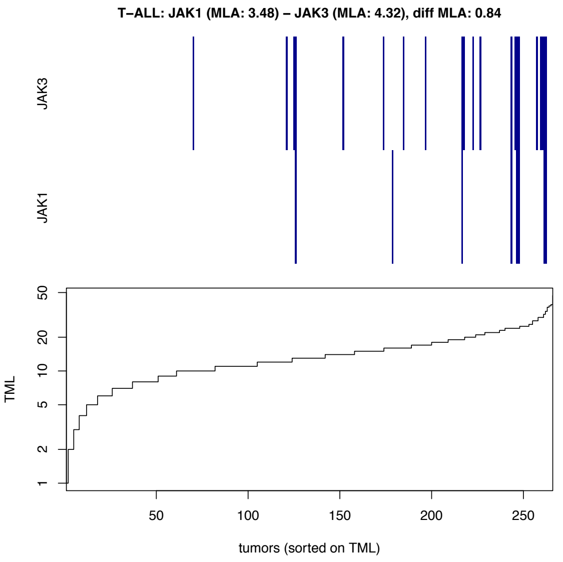
T–ALL: JAK1 (MLA: 3.48) – JAK3 (MLA: 4.32), diff MLA: 0.84
JAK3
JAK1
TML
1
2
5
10
20
50
50
100
150
200
250
tumors (sorted on TML)

G

***Supplemental Figure S4: Comparison p-values Permutation and WeSME test***

P-values resulting from testing gene pairs in all cancer types (excluding the PAN cancer test) with the Permutation test compared to their corresponding p-values produced with the WeSME test. The WeSME test was run ten times, but only the results from the first run are shown for better comparison. Each dot represents a gene pair tested for co-occurrence (CO, blue) or mutual exclusivity (ME, red). Only gene pairs with co-occurrence count > 2 were tested (see Materials and Methods), explaining the small proportion of CO data points.

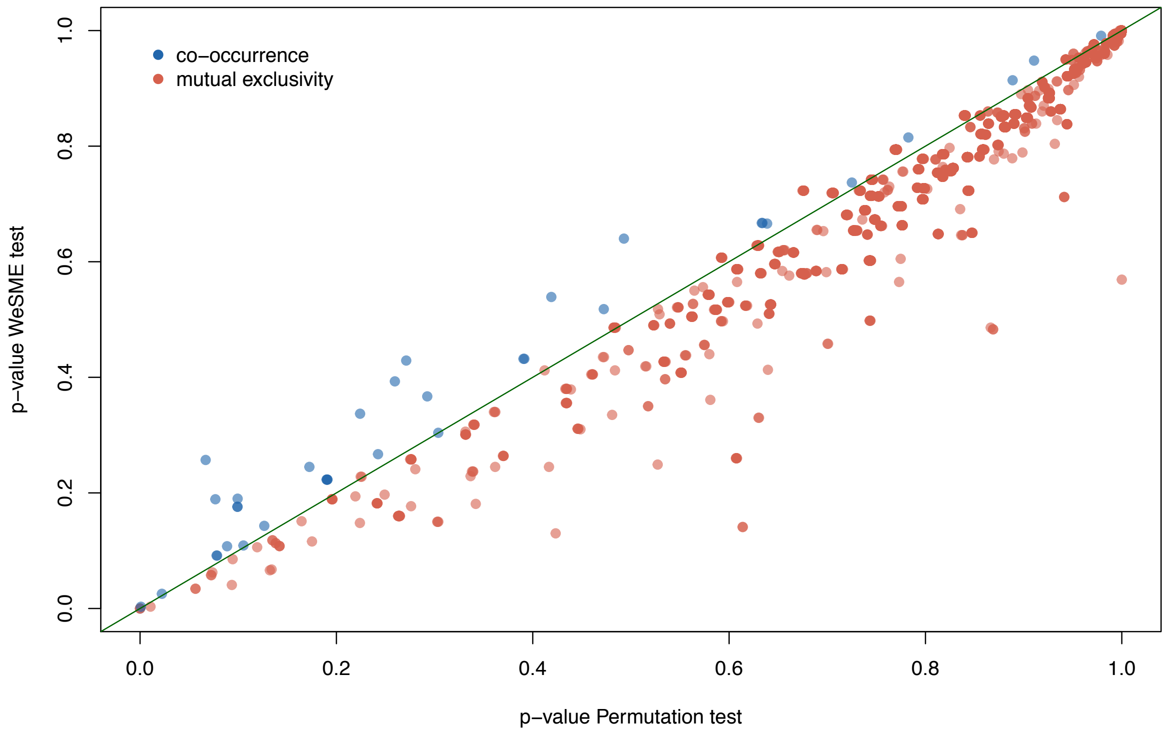

*Supplemental Figure S5: Detailed workflow of Genetic Interactions pipeline*

Starting with a gene-sample mutation matrix, the permutation tests counts the number of co-occurrences for each gene pair and compares this count to a null distribution based on matrix randomizations (*N=1,000,000*) to compute p-values. The same gene-sample matrix serves as input for the WeSME test. This test compares mutual exclusivity counts with a null distribution generated by weighted sampling based on sample-level mutation rates. False discovery rates are estimated by comparing p-values with a null distribution of p-values. This null-distribution is created by testing 100 (Permutation test) or 300 (WeSME test) random matrices. This figure is based on Figure 1 in Park and Lehner[23] and Figure 1 in Kim, Madan, and Przytycka[28].

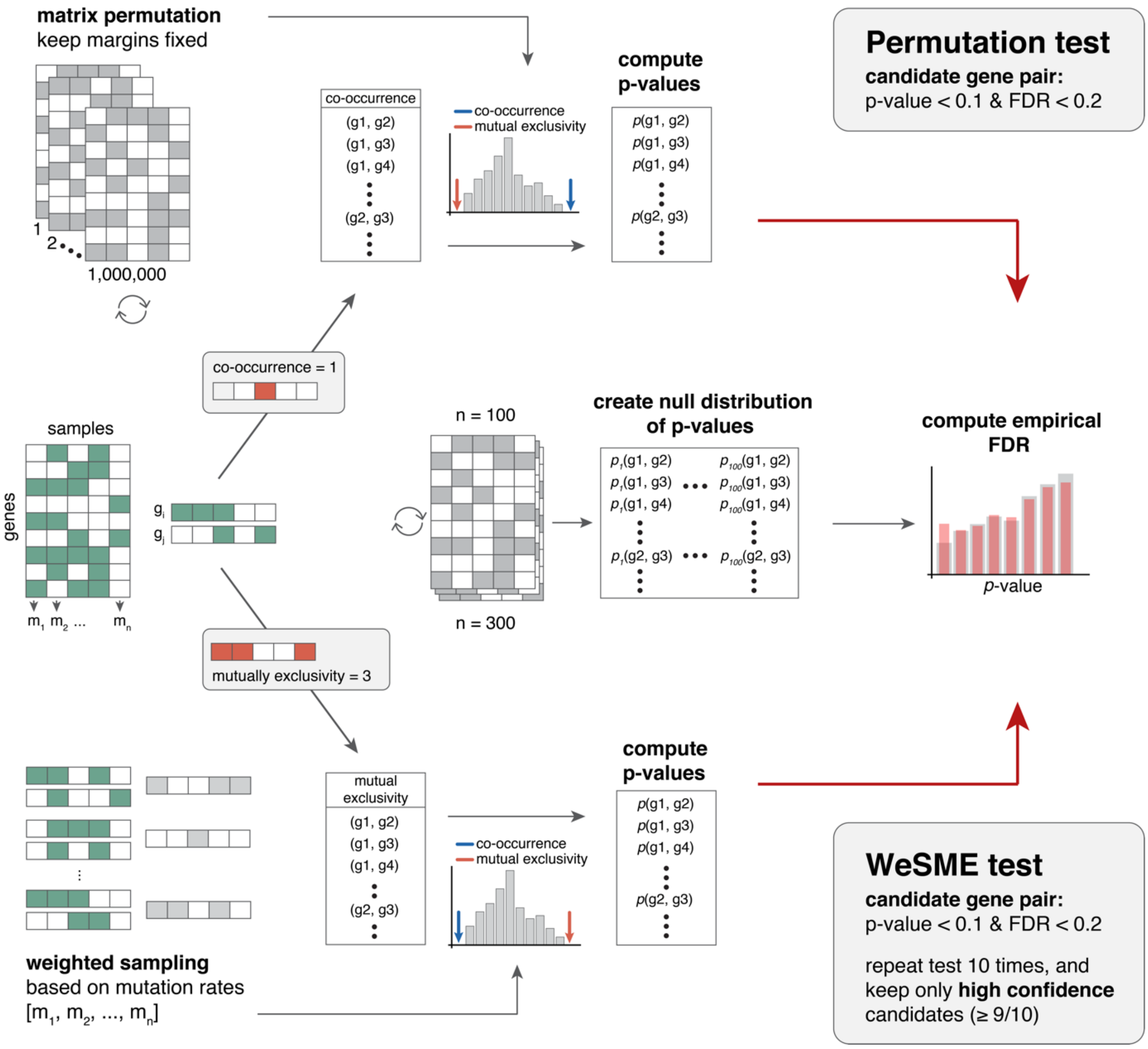
